# Anemonefish are better taxonomists than humans

**DOI:** 10.1101/2023.09.11.557259

**Authors:** Rio Kashimoto, Manon Mercader, Jann Zwahlen, Saori Miura, Miyako Tanimoto, Kensuke Yanagi, James Davis Reimer, Konstantin Khalturin, Vincent Laudet

**Affiliations:** Marine Eco-Evo-Devo Unit, Okinawa Institute of Science and Technology. 1919-1 Tancha, Onna-son, Okinawa 904-0495, Japan; Marine Genomics Unit, Okinawa Institute of Science and Technology. 1919-1 Tancha, Onna-son, Okinawa 904-0495, Japan; Okinawa Churaumi Aquarium, 424 Ishikawa, Motobu, Kunigami District, Okinawa 905-0206, Japan; Coastal Branch of Natural History Museum and Institute, Chiba, Katsuura, Chiba, Japan; Graduate School of Engineering and Science, University of the Ryukyus, 1 Senbaru, Nishihara, Okinawa 903-0213, Japan; Tropical Biosphere Research Center, University of the Ryukyus, 1 Senbaru, Nishihara, Okinawa 903-0213, Japan; Institute of Cellular and Organismic Biology (ICOB), Academia Sinica, 128 Academia Road, Section 2, Nangang, Taipei 115, Taiwan; Marine Research Station, Institute of Cellular and Organismic Biology (ICOB), Academia Sinica, 23-10, Dah-Uen Rd, Jiau Shi, I-Lan 262, Taiwan

## Abstract

The symbiosis between giant sea anemones, photosynthetic algae of the family Symbiodiniaceae, and anemonefish is an iconic example of a mutualistic “menage à 3”^1^. Patterns of associations among 28 species of anemonefish and 10 species of giant sea anemone hosts are complex: Some anemonefish species are highly specialized to inhabit only one species of sea anemone (*e*.*g*., *Amphiprion frenatus* with *Entacmaea quadricolor*), whereas others are more generalist and can live in almost any host species (e.g., *Amphiprion clarkii*)^1,2,3^. Reasons for host preferences and the mechanisms involved are obscured, among other things, by the lack of resolution of giant sea anemone phylogeny. Recent molecular analyses have shown that giant sea anemones hosting anemonefish belong to three distinct clades: *Entacmaea, Stichodactyla*, and *Heteractis*^4,5,6^. Inside these clades, however, species delimitation has been impeded by morphological variability of the giant sea anemones and is poorly resolved with classical markers^4,5^. Here, we employed an extensive transcriptomic dataset from 55 sea anemones collected from southern Japan to build a robust phylogeny. With this dataset, we observed that the bubble-tip sea anemone *E. quadricolor*, currently considered to be a single species, can in fact be separated into at least four distinct cryptic lineages (A-D). Moreover, these lineages can be precisely distinguished by their association with anemonefish: *A. frenatus* is only found associated with lineage D, whereas *A. clarkii* lives in the other three lineages.

---

We built a robust phylogeny with a transcriptomic dataset of 55 samples of sea anemones collected across the Ryukyu Archipelago to mainland Japan as well as from the isolated Ogasawara Islands (See Sup. Table; Sup Fig 1A). RNA was extracted from tentacles and on average 65.3 million paired-end 150bp reads per samples were sequenced on a NovSeq6000. Data processing and filtering, including the removal of Symbiodiniaceae sequences, was performed as previously described^6^. We generated a collection of 55 high quality transcriptomes that were used to build phylogenetic trees^7^ and analyze gene expression variation among the individuals.

**Figure legend.**
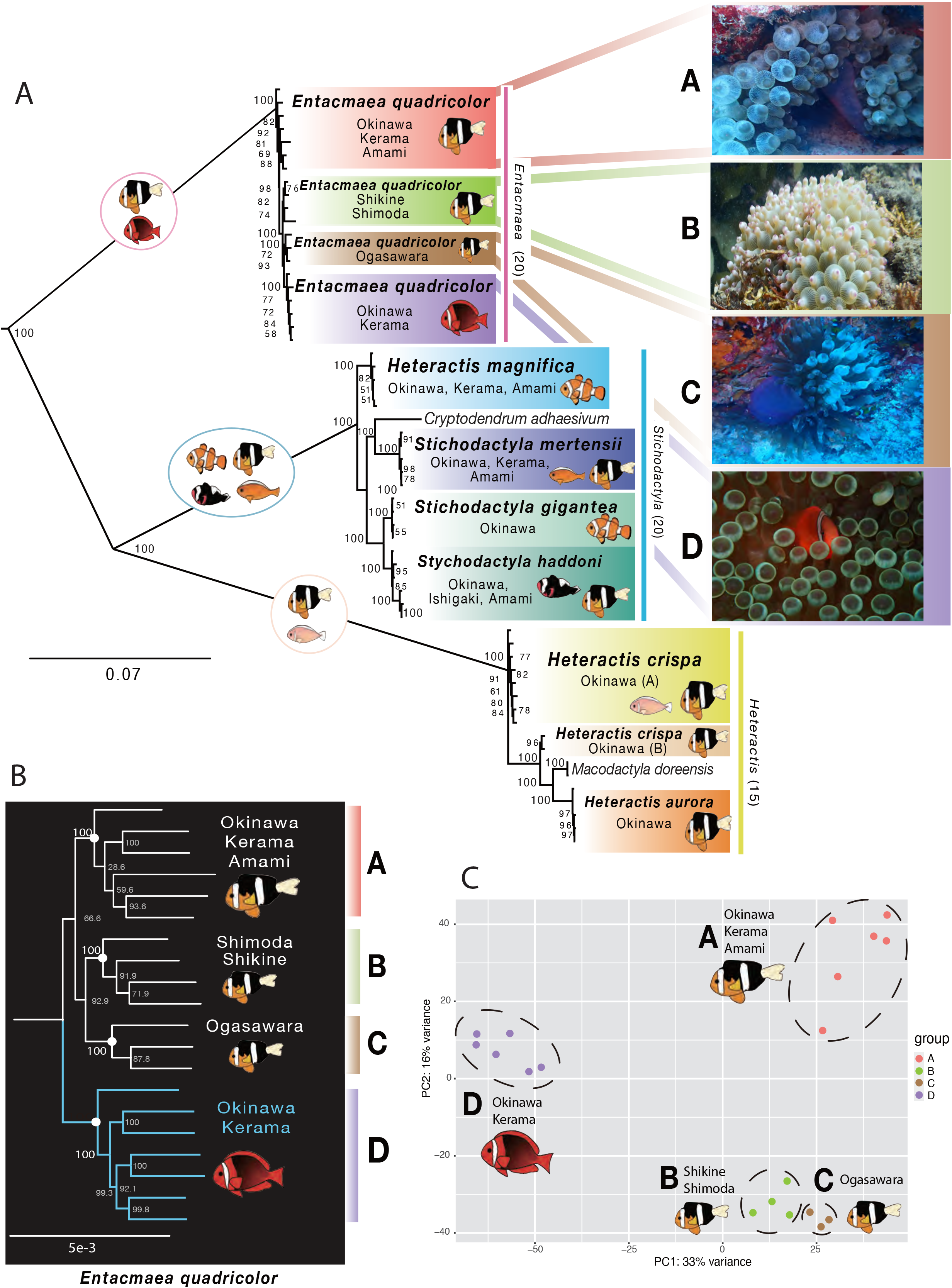
(A) ML topology obtained using the supermatrix and a best-fit partitioning scheme in IQ-TREE (Nguyen et al., 2015). Numbers in parentheses in the three main clades (*Entacmaea, Stichodactyla, Heteractis*) are numbers of samples in our dataset for each clade. (B) Detailed topology focusing on *E. quadricolor* clade based on a dataset containing 1122 genes. The four lineages are monophyletic and supported by high bootstrap values. (C) Principal Component Analysis of gene expression showing PC1 axis differentiation according to anemonefish species and PC2 according to geographical position.

Initial phylogenetic analysis relied on a supermatrix composed of 28,608 amino acid sites from 110 genes and included datasets from NCBI (*Cryptodendrum adhaesivum, Macrodactyla doreensis*), with *Exaiptasia* used as outgroup (Fig. 1A). Inference was performed using Maximum Likelihood^7^. While recovering a topology similar to those of previous studies^4,5,6^, our analysis was the first to clearly resolve putative species boundaries among the 3 major clades: *Heteractis* (two clusters of *Heteractis crispa*, as well as one of *M. doreensis* and *H. aurora*), *Stichodactyla* (*S. mertensii, S. gigantea, S. haddoni, H. magnifica, C. adhaesivum*), and *Entacmaea*.

Interestingly, we observed that *Entacmaea* was divided into 4 lineages, and we therefore conducted more focused phylogenetic analyses with larger gene sets (Figure 1B, Sup Fig 1B). We observed monophyletic lineages: A, that contained specimens from Okinawa, Kerama, and Amami Islands, B, from Shikine island and Shimoda; C, from the Ogasawara Islands, and D, from Okinawa and Kerama Islands. While the differentiation between lineages A, B, and C may be classical geographical allopatry, we observed that lineages A and D lived in sympatry but hosted different anemone fish species: *A. clarkii* for lineage A and *A. frenatus* for lineage D. This association between anemone fish species and host anemone lineage was highly significant (Fisher’s exact test, P= 3.45E^-08^). Topology was remarkably stable, with most internal nodes fully supported across all analyses using various gene sets and outgroups (Sup Fig 1B). The overall genetic distance between lineage A and D was in the same range but slightly lower than that observed between different species of *Stichodactyla* (Fig 1A). Our phylogenetic tree (Fig. 1B) suggested that *Entacmaea* first split between an *A. clarkii-*hosting branch (lineages A, B, and C) and an *A. frenatus*-hosting one (lineage D). Given the relatively low levels of genetic distance between lineages A and D and their sympatric distribution, we may be observing the process of species differentiation between these lineages. To further ascertain the validity of this association, we conducted choice experiments in aquaria, in which either *A. clarkii* or *A. frenatus* naïve juveniles were given a choice between lineage A or lineage D of *Entacmaea* (Sup Fig 2A). We observed that no *A. clarkii* chose lineage D whereas *A. frenatus* mostly, but not always, chose lineage D (Sup Fig 2B). Of note, some fish (6/20 *A. clarkii* and 7/20 *A. frenatus*) did not exhibit a clear choice. These data show that even in captivity, naïve juveniles of these two species significantly (Fisher’s exact test, P= 5.98E^-06^) reproduced the association patterns that we detected in the wild, despite having never encountered sea anemones before.

Genetic differences revealed by phylogenetic analyses are paralleled by clear differences in global gene expression among the 4 clades of *Entacmaea* (Figure 1C). The largest observed variation was associated with the anemonefish hosts (*A. frenatus* vs. *A. clarkii*) whereas the second component separated lineages A, B, and C according to geographic location: northern samples being widely separated from southern ones. We found 3699 differentially expressed genes (DEGs) with 4-fold differences in expression among all 4 lineages (Sup Fig. 3A,B), and 1397 DEGs between the two sympatric lineages A and D (Sup Fig. 3C,D). Among these, 513 encoded predicted proteins, many of which (427) are likely cnidarian-specific genes of unknown function. Interestingly, among the DEGs among the clades were fluorescent proteins^8^ (Sup Fig.2E) and putative nematocyte-specific genes (Sup Fig.3F). To human eyes clades A and D do not have any noticeable coloration difference, but clownfish might have different perception. Contribution of nematocyte-specific genes is easier to interpret since it is known that stinging capacity differs among the species of sea anemones with *Stichodactyla* being the most toxic^9^. It might be that the clades A and D of *Entacmaea* have acquired differences in stinging capacity which make them more or less preferable hosts for *A. clarkii* and *A. frenatus*. Cause and effect relationships require further investigations, but our analysis suggests that ongoing speciation within *Entacmaea* is already accompanied by considerable differences in gene expression.

Now it will be important to clarify whether any morphological synapomorphies are associated with the various lineages and how these impact the symbioses^10^. Also, it must be determined who is the driver of the detected divergences: is it the anemonefish that induces speciation or is it sea anemone that adapts to different species of fish? Also, our results implies that the classical table^1,3^ describing the specific relationships between the anemonefish and their host should be revisited. It is striking that these two fish species can recognize distinct lineages that taxonomists have not been able to clearly separate until now. In this sense, anemonefish appear to be better ‘taxonomists’ than humans.

## Acknowledgements

We thank Nori Satoh for constant support and advice, Felipe Castellano-Macias for help in the statistical analyses, Kina Hayashi for the anemonefish drawings, and Marcela Herrera and Agneesh Barua for critical reading of this manuscript. This research has been supported by a KICKS OIST grant to V.L. and J.D.R.

---

**Figure S1**

(A) Sampling locations in the southern and central Japan are shown with filled arrowheads. Detailed information is provided in Table 1. (B) Phylogenetic reconstructions obtained with concatenated alignments of 1006, 1107 and 1122 genes in IQ-TREE. Four clades within the genus *Entacmaea* with high bootstrap support were revealed in all the analyses. Color bars are used to label samples belonging to clades A, B, C and D.

**Figure S2**

(A) Histogram depicting the selection of lineages A and D by *A. clarkii* and *A. frenatus* juveniles in a choice experiment. (B) Choice experiments were led in a 130x67x40 cm plastic tank. One *E. quadricolor* of lineage A and one of lineage D were placed in a plastic basket at each side of the tank to prevent them from moving away. They were maintained with a constant flow of natural seawater which was turned off when performing the experiment. At the start of each trial, the sea anemones’ diameter was measured to account for biotic variations throughout the experimental period. A juvenile fish (2-5cm) was introduced halfway between the two anemones in a little box with a tight grid. Each individual was allowed to recover for 5 minutes before the box/basket was carefully lifted to release into the experimental arena for an experimental duration of 30 minutes, which was recorded by cameras (GoPro Hero 7 or 8) placed on the long sides of the tank. The fish was considered to be making a choice if entering one of the anemones by swimming through the tentacles and staying associated with it for the remainder of the experiment. Fish swimming around, staying in the middle of the tank or even on one side, or in/under the basket were considered not selecting a host. Twenty trials were done for each fish species (*A. clarkii* and *A. frenatus*). The anemones were switched sides regularly to avoid any bias due to potential side preference, an equal number of trials were done for each side.

**Figure S3**

(A) Heatmap shows the expression of 3699 genes that are differentially expressed among the samples representing clades A,B,C,D (4-fold difference cut-off, P=0.01). (B) Sample correlation heatmap and clustering of samples based on gene expression. Sub-set of *Entacmaea* samples was used. (C) Heatmap shows expression of 1397 genes that are differentially expressed between the samples belonging to clades A and D (4-fold difference cut-off, P=0.01). (D) Sample correlation heatmap and clustering of samples based on gene expression. Samples belonging to clades A and D were compared. (E) Expression of all genes that encode putative fluorescent proteins in 20 samples of *Entacmaea*. Samples are grouped by their respective clades (D,A,B,C) and the genes were clustered based on their expression levels. Log2 transformed TMM normalized FPKM values were used. (F) Expression of putative nematocyte-specific genes with 4-fold expression difference between clades A and D. Genes were identified by BLAST search against the reference set of *Hydra* nematocyte-specific proteins with E-value cut-off 1e-5. Gene identifiers (transcript IDs) and well as protein annotations are shown at the right side of the heatmaps in E and F. Sample names are shown at the bottom of the heatmaps. Order of samples is identical in A,B and C,D and E,F. Detailed information about the samples is shown in Table 1.

## References

1. Hoepner, C.M., Fobert, E.K., Abbott, C.A., and Burke da Silva, K. (2022) No place like home: can omics uncover the secret behind the sea anemone and anemonefish symbiotic relationship. in Laudet V and Ravasi T “Evolution Development and Ecology of Anemonefish. Model organisms for Marine Science” CRC Press. Boca-Raton pp. 306.

2. Roux, N., Salis, P., Lee, S.H., Besseau, L. and Laudet, V. (2020) Anemone fishes, a model for Eco-Evo-Devo. EvoDevo, 11, 20. 10.1186/s13227-020-00166-7

3. Fautin, D.G. (1991). The anemonefish symbiosis: what is known and what is not. Symbiosis 10, 23–46. http://hdl.handle.net/1808/6134

4. Titus, B.M., Benedict, C., Laroche, C., Gusmão, L.C., Van Deusen, V., Chiodo, T., Meyer, C.P., Berumen, M.L., Bartholomew, A., Yanagi, K., Reimer, J.D., Fujii, T., Daly and M., Rodríguez, E. (2019) Phylogenetic relationships among the clownfish-hosting sea anemones. Mol. Phyl. Evol. 139, 106526. 10.1016/j.ympev.2019.106526

5. Nguyen, H.T.T., Dang, B.T., Glenner, H. and Geffen A.J. (2019). Cophylogenetic analysis of the relationship between anemonefish Amphiprion (Perciformes: Pomacentridae) and their symbiotic host anemones (Anthozoa: Actiniaria). Mar. Biol. Res. 16, 117–133. 10.1080/17451000.2020.1711952

6. Kashimoto, R., Tanimoto, M., Miura, S., Satoh, N., Laudet, V. and Khalturin, K. (2022) Transcriptomes of giant sea anemones from Okinawa as a tool for understanding their phylogeny and symbiotic relationships with anemonefish. Zool. Sci. 39, 374–387. 10.2108/zs210111

7. Zhang, C., Rabiee, M., Sayyari, E., and Mirarab, S. (2018). ASTRAL-III: polynomial time species tree reconstruction from partially resolved gene trees. BMC Bioinformatics 19, 153. 10.1186/s12859-018-2129-y

8. Kashimoto, R., Hisata, K., Shinzato, C., Satoh, N., and Shoguchi, E. (2021). Expansion and diversification of fuorescent protein genes in fifteen Acropora species during the evolution of acroporid corals. Genes 12, 397. 10.3390/genes12030397

9. Nedosyko, A.M., Young, J.E., Edwards, J.W., Burke da Silva K. (2014) Searching for a toxic key to unlock the mystery of anemonefish and anemone symbiosis. PLoS One 9, e98449. 10.1371/journal.pone.0098449

10. Ashwood, L.M., Mitchell, M.L., Madio, B., Hurwood, D.A., King, G.F., Undheim, E.A.B., Norton, R.S., and Prentis, P.J. (2021). Tentacle morphological variation coincides with differential expression of toxins in sea anemones. Toxins 13, 452. 10.3390/toxins13070452

