## Supplemental Figure for "Anemonefish are better taxonomists than humans": KashimotoBioRxiv (dragged) 2.pdf

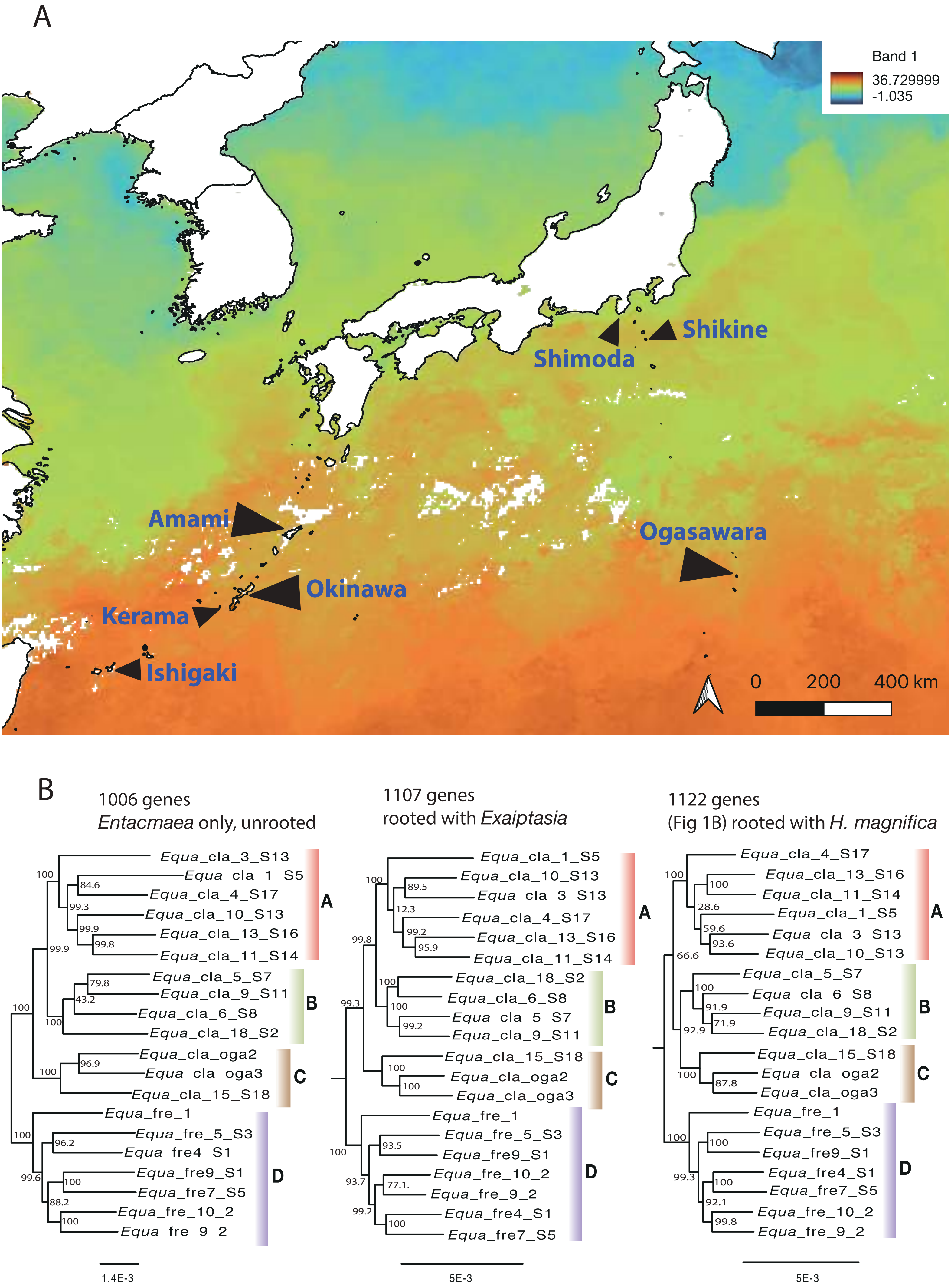


Kashimoto et al., Figure S1

A

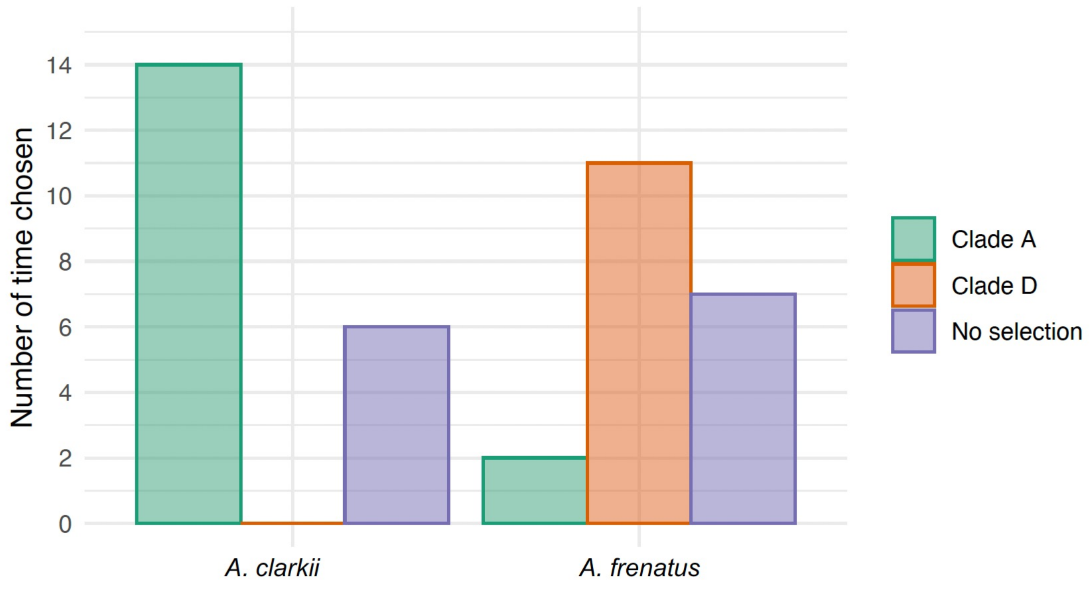


B

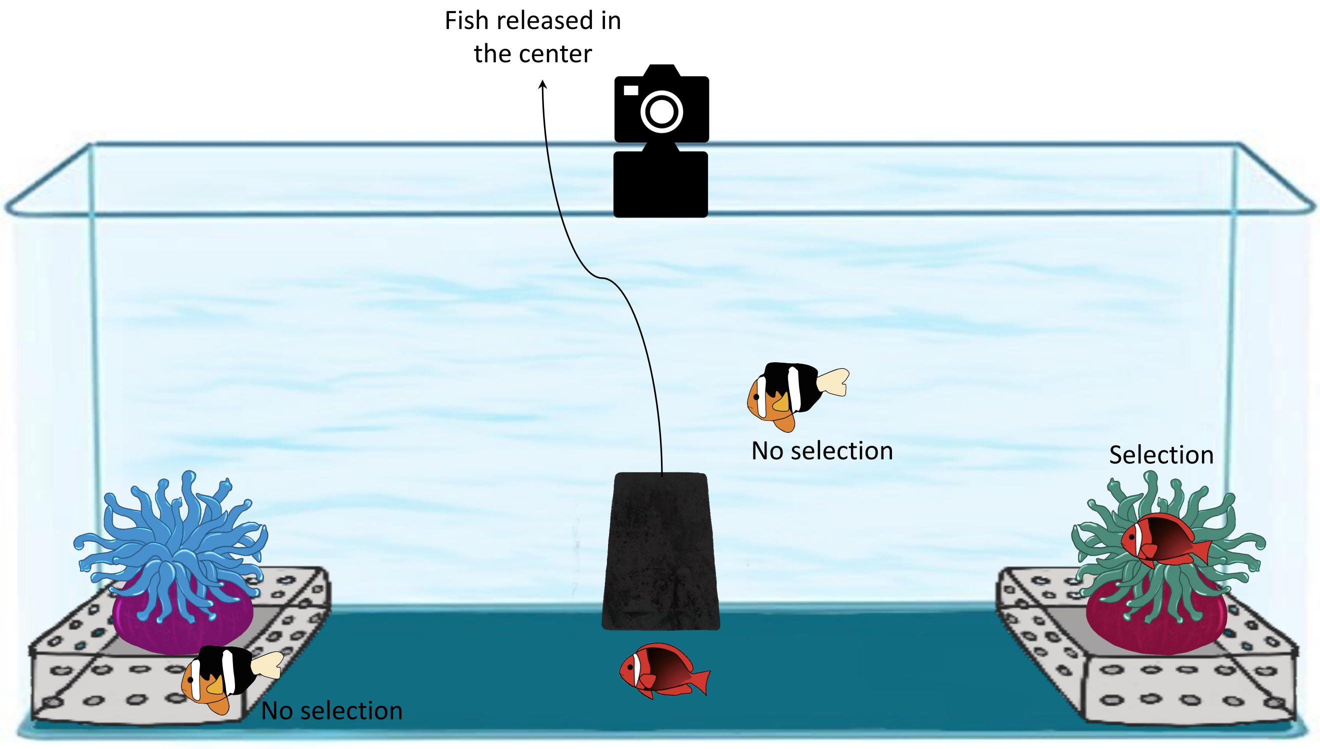


SupFigure2

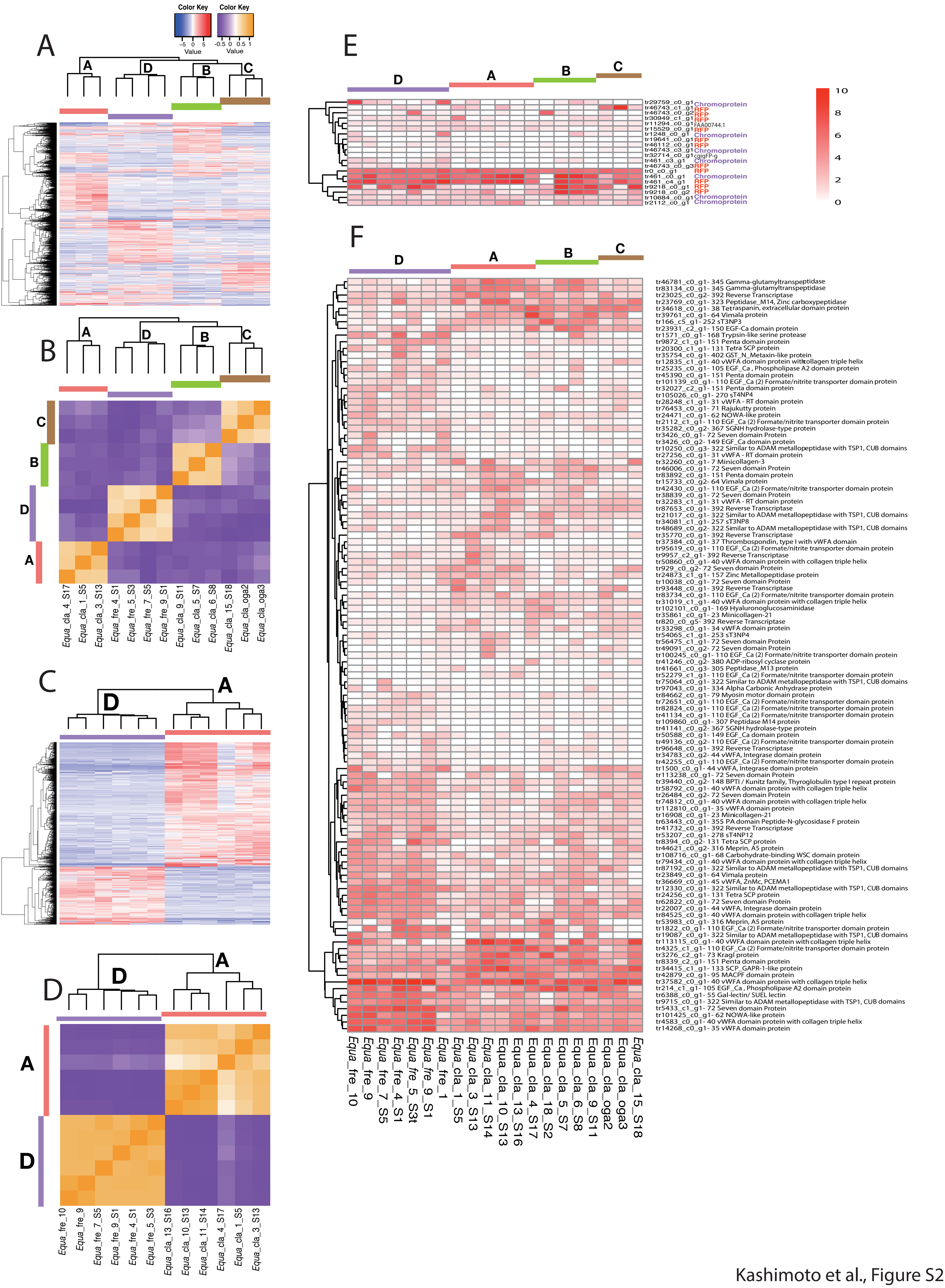


Kashimoto et al., Figure S2

| Sample No | Sample ID | Species | Hosting anemonefish | Second hosting anemonefish | Location | SRA | Data amount per species (million reads) | Yield (Mbases) | Total Transcripts | N50 | Longest Transcript | BUSCO (transcriptome) | GC% |
|---|---|---|---|---|---|---|---|---|---|---|---|---|---|
| 1 | Equa_cla_1_S5 | *Entacmaea quadricolor* | *Amphiprion clarkii* | | Kerama island, Okinawa | SRR23908452 | 56.9 | 8,588 | 189,103 | 890 | 24,489 | C:77.0%[S:48.5%,D:28.5%],F:12.6%,M:10.4%,n:954 | 37.54 |
| 2 | Equa_cla_3_S13 | *Entacmaea quadricolor* | *Amphiprion clarkii* | | Urasoe, Okinawa | SRR21497999 | 52.2 | 7,877 | 135,764 | 847 | 20,618 | C:70.1%[S:50.2%,D:19.9%],F:16.5%,M:13.4%,n:954 | 37.54 |
| 3 | Equa_cla_4_S17 | *Entacmaea quadricolor* | *Amphiprion clarkii* | | Onna, Okinawa | SRR23908441 | 55.0 | 8,308 | 115,104 | 1,072 | 17,027 | C:82.6%[S:64.8%,D:17.8%],F:6.1%,M:11.3%,n:954 | 37.47 |
| 4 | Equa_cla_10_S13 | *Entacmaea quadricolor* | *Amphiprion clarkii* | | Amami, Kagoshima | SRR21498014 | 46.4 | 7,011 | 83,140 | 1,465 | 32178 | C:78.7%[S:59.7%,D:19.0%],F:8.6%,M:12.7%,n:954 | 38.26 |
| 5 | Equa_cla_11_S14 | *Entacmaea quadricolor* | *Amphiprion clarkii* | | Amami, Kagoshima | SRR21498013 | 50.3 | 7,599 | 79,734 | 1,531 | 31722 | C:81.1%[S:65.3%,D:15.8%],F:7.2%,M:11.7%,n:954 | 38.44 |
| 6 | Equa_cla_13_S16 | *Entacmaea quadricolor* | *Amphiprion clarkii* | | Amami, Kagoshima | SRR23908453 | 55.3 | 8,357 | 72,257 | 1,415 | 22968 | C:75.3%[S:59.9%,D:15.4%],F:10.6%,M:14.1%,n:954 | 38.47 |
| 7 | Equa_cla_18_S2 | *Entacmaea quadricolor* | *Amphiprion clarkii* | | Shikine-jima, Tokyo | SRR21498002 | 31.1 | 4,700 | 127,636 | 1,267 | 21646 | C:85.1%[S:57.0%,D:28.1%],F:6.3%,M:8.6%,n:954 | 37.53 |
| 8 | Equa_cla_5_S7 | *Entacmaea quadricolor* | | Dascylus trimaculatus | Shimoda, Shizuoka | SRR23908437 | 53.9 | 8,144 | 78,100 | 1,468 | 21941 | C:81.9%[S:62.6%,D:19.3%],F:7.7%,M:10.4%,n:954 | 38.54 |
| 9 | Equa_cla_6_S8 | *Entacmaea quadricolor* | | Dascylus trimaculatus | Shimoda, Shizuoka | SRR23908436 | 46.1 | 6,962 | 83,143 | 1,326 | 31464 | C:76.1%[S:59.9%,D:16.2%],F:9.6%,M:14.3%,n:954 | 38.78 |
| 10 | Equa_cla_9_S11 | *Entacmaea quadricolor* | | | Shikine-jima, Tokyo | SRR23908435 | 41.6 | 6,282 | 59,911 | 1,312 | 31703 | C:66.5%[S:54.6%,D:11.9%],F:14.8%,M:18.7%,n:954 | 39.04 |
| 11 | Equa_cla_15_S18 | *Entacmaea quadricolor* | *Amphiprion clarkii* | | Ogasawara, Tokyo | SRR23908442 | 45.1 | 6,806 | 63,554 | 1,364 | 26407 | C:62.5%[S:50.7%,D:11.8%],F:17.6%,M:19.9%,n:954 | 38.7 |
| 12 | Equa_cla_oga2 | *Entacmaea quadoricolor* | *Amphiprioan clarkii* | | Ogasawara, Tokyo | SRR23908451 | 50.1 | 7,559 | 97,830 | 1712 | 32449 | C:81.2%[S:60.2%,D:21.0%],F:7.5%,M:11.3%,n:954 | 44.98 |
| 13 | Equa_cla_oga3 | *Entacmaea quadoricolor* | *Amphiprioan clarkii* | | Ogasawara, Tokyo | SRR23908450 | 50.8 | 7,676 | 107,232 | 1676 | 32901 | C:90.6%[S:62.6%,D:28.0%],F:1.9%,M:7.5%,n:954 | 43.09 |
| 14 | Equa_fre_4_S1 | *Entacmaea quadricolor* | *Amphiprion frenatus* | | Aka-jima, Okinawa | SRR23908434 | 40.7 | 6,146 | 67,738 | 1,275 | 17,105 | C:67.4%[S:52.9%,D:14.5%],F:18.4%,M:14.2%,n:255 | 39.15 |
| 15 | Equa_fre_5_S3 | *Entacmaea quadricolor* | *Amphiprion frenatus* | | Aka-jima, Okinawa | SRR23908433 | 53.7 | 8,108 | 66,148 | 1,211 | 18,074 | C:63.6%[S:50.6%,D:13.0%],F:19.0%,M:17.4%,n:954 | 39.32 |
| 16 | Equa_fre_9_S1 | *Entacmaea quadricolor* | *Amphiprion frenatus* | | Kerama island, Okinawa | SRR23908431 | 49.5 | 7,479 | 100,951 | 1,578 | 32953 | C:88.6%[S:64.2%,D:24.4%],F:3.4%,M:8.0%,n:954 | 38.05 |
| 17 | Equa_fre_7_S5 | *Entacmaea quadricolor* | Amphiprion frenatus | | Aka-jima, Okinawa | SRR23908432 | 49.5 | 7,479 | 61,916 | 1,373 | 19904 | C:70.0%[S:56.1%,D:13.9%],F:13.8%,M:16.2%,n:954 | 39.34 |
| 18 | Equa_fre_9_2 | *Entacmaea quadricolor* | *Amphiprion frenatus* | | Yomitan, Okinawa | SRR14307520 | 42.9 | 5,660 | 134,868 | 1,561 | 20,945 | C:78.7%[S:54.4%,D:24.3%],F:8.3%,M:13.0%,n:954 | 46.2 |
| 19 | Equa_fre_10_2 | *Entacmaea quadricolor* | *Amphiprion frenatus* | | Yomitan, Okinawa | SRR14307530 | 37.5 | 6,478 | 127,976 | 1,526 | 19233 | C:75.5%[S:55.9%,D:19.6%],F:11.7%,M:12.8%,n:954 | 45.8 |
| 20 | Equa_fre_1 | *Entacmaea quadricolor* | *Amphiprion frenatus* | | Okinawa (Aquarium) | SRR14307527 | 145 | 21,910 | 341,093 | 1,848 | 33,093 | C:90.5%[S:50.7%,D:39.8%],F:3.7%,M:5.8%,n:954 | 43.3 |
| 21 | Hmag4 | *Heteractis magnifica* | *Amphiprion ocellaris* | | Onna, Okinawa | SRR21498011 | 59.0 | 8,905 | 89,926 | 1,451 | 25271 | C:83.7%[S:58.1%,D:25.6%],F:7.0%,M:9.3%,n:954 | 39.56 |
| 22 | Hmag_5 | *Heteractis magnifica* | *Amphiprion ocellaris* | | Yomitan, Okinawa | SRR21498010 | 54.3 | 8,198 | 89,927 | 1,195 | 12535 | C:71.0%[S:48.4%,D:22.6%],F:13.9%,M:15.1%,n:954 | 39.19 |
| 23 | Hmag_3_2_S20 | *Heteractis magnifica* | *Amphiprion ocellaris* | | Naze, Kogoshima | SRR21498012 | 55.5 | 8,379 | 121,881 | 1,694 | 26502 | C:91.2%[S:58.1%,D:33.1%],F:2.7%,M:6.1%,n:954 | 39.13 |
| 24 | Hmag_1 | *Heteractis magnifica* | *Amphiprion ocellaris* | | Ginowan, Okinawa | SRR14307526 | 148.9 | 22,489 | 286,530 | 2,045 | 35,951 | C:92.5%[S:53.4%,D:39.1%],F:2.4%,M:5.1%,n:954 | 44.3 |
| 25 | Hmag_22 | *Heteractis magnifica* | *Amphiprion ocellaris* | | Yomitan, Okinawa | SRR14307529 | 34.5 | 6945 | 121,458 | 1,413 | 26376 | C:89.7%[S:56.1%,D:33.6%],F:3.2%,M:7.1%,n:954 | 45 |
| 26 | Smer_2_S8 | *Stichodactyla mertensii* | *Amphiprion clarkii* | | Kerama island, Okinawa | SRR21498005 | 50.4 | 7,615 | 86,778 | 1556 | 19163 | C:86.7%[S:58.1%,D:28.6%],F:5.6%,M:7.7%,n:954 | 40.02 |
| 27 | Smer_3_S9 | *Stichodactyla mertensii* | *Amphiprion clarkii* | *Amphiprion sandaracinos* | Kerama island, Okinawa | SRR21498004 | 48.6 | 7,341 | 91,846 | 1,788 | 32913 | C:89.9%[S:59.3%,D:30.6%],F:3.1%,M:7.0%,n:954 | 39.56 |
| 28 | Smer_4_S16 | *Stichodactyla mertensii* | *Amphiprion clarkii* | *Amphiprion sandaracinos* | Tancha, Okinawa | SRR21498003 | 60.1 | 9,080 | 127,235 | 1,284 | 30046 | C:81.3%[S:53.4%,D:27.9%],F:10.0%,M:8.7%,n:954 | 39.13 |
| 29 | Smer_6_S12 | *Stichodactyla mertensii* | *Amphiprion clarkii* | Amphiprioan sandaracinos | Setouchi, Kagoshima | SRR21498001 | 39.8 | 6,016 | 89,171 | 1,633 | 34462 | C:84.0%[S:63.4%,D:20.6%],F:6.8%,M:9.2%,n:954 | 39.25 |
| 30 | Smer_1 | *Stichodactyla mertensii* | *Amphiprion polymnus* | | Okinawa (aquerium) | SRR14307525 | 156.2 | 23,580 | 269,130 | 2,063 | 37,745 | C:91.9%[S:56.6%,D:35.3%],F:2.4%,M:5.7%,n:954 | 45.5 |
| 31 | Sgig_3 | *Stichodactyla gigantea* | *Amphiprion ocellaris* | | Onna, Okinawa | SRR21498009 | 55.5 | 8,379 | 97,011 | 1,692 | 33169 | C:86.7%[S:57.9%,D:28.8%],F:6.0%,M:7.3%,n:954 | 39.61 |
| 32 | Sgig_4 | *Stichodactyla gigantea* | *Amphiprion ocellaris* | | Onna, Okinawa | SRR21498008 | 56.3 | 8,500 | 113,582 | 1,671 | 32577 | C:90.5%[S:55.9%,D:34.6%],F:3.5%,M:6.0%,n:954 | 39.41 |
| 33 | Sgig_1 | *Stichodactyla gigantea* | *Amphiprion ocellaris* | | Okinawa (Aquerium) | SRR14307524 | 169.6 | 25,612 | 323,507 | 2,178 | 64,912 | C:92.5%[S:47.0%,D:45.5%],F:2.0%,M:5.5%,n:954 | 43.7 |
| 34 | Sgig_2 | *Stichodactyla gigantea* | *Amphiprion ocellaris* | | Onna, Okinawa+G37 | SRR14307528 | 46 | 6208 | 118,439 | 1,516 | 24056 | C:80.9%[S:55.3%,D:25.6%],F:8.5%,M:10.6%,n:954 | 45 |
| 35 | Shad_2 | *Stichodactyla haddoni* | *Amphiprion polymnus* | | Seragaki, Okinawa | SRR21498007 | 60.8 | 9,174 | 93,837 | 1,685 | 31815 | C:88.1%[S:64.0%,D:24.1%],F:4.3%,M:7.6%,n:954 | 39.64 |
| 36 | Shad_3 | *Stichodactyla haddoni* | *Amphiprion polymnus* | | Seragaki, Okinawa | SRR21498006 | 63.3 | 9,561 | 93,837 | 1,748 | 32056 | C:84.2%[S:60.8%,D:23.4%],F:6.8%,M:9.0%,n:954 | 39.79 |
| 37 | Shad_1 | *Stichodactyla haddoni* | no fish | | Okinawa (aquerium) | SRR14307523 | 186.5 | 28,156 | 343,778 | 2,302 | 53,044 | C:92.7%[S:44.8%,D:47.9%],F:2.1%,M:5.2%,n:954 | 43.8 |
| 38 | Shad_5_S20 | *Stichodactyla haddoni* | *Amphiprioan clarkii* | | Ishigaki, Okinawa | SRR23908440 | 47.7 | 7199 | 153131 | 1933 | 32079 | C:90.7%[S:62.6%,D:28.1%],F:3.0%,M:6.3%,n:954 | 44.59 |
| 39 | Shad_12_S15 | *Stichodactyla haddoni* | *Amphiprioan clarkii* | | Setouch, Kogoshima | SRR23908438 | 48.9 | 7,388 | 96469 | 1759 | 32175 | C:90.2%[S:59.1%,D:31.1%],F:3.4%,M:6.4%,n:954 | 39.56 |
| 40 | Shad_6 | *Stichodactyla haddoni* | *Amphiprioan clarkii* | | Tancha, Okinawa | SRR23908439 | 42.1 | 6353 | 125355 | 1931 | 33007 | C:85.4%[S:64.0%,D:21.4%],F:5.2%,M:9.4%,n:954 | 45.51 |
| 41 | Hcri_3_S2 | *Heteractis crispa* | *Amphiprion perderaion* | | Akajima, Okinawa | SRR23908448 | 42.7 | 6,450 | 68,293 | 1,563 | 26,495 | C:81.3%[S:59.2%,D:22.1%],F:8.7%,M:10.0%,n:954 | 41.84 |
| 42 | Hcri_4_S4 | *Heteractis crispa* | *Amphiprion perderaion* | | Kerama island, Okinawa | SRR21497994 | 51.1 | 7,715 | 59,769 | 1,604 | 27485 | C:80.1%[S:62.4%,D:17.7%],F:8.6%,M:11.3%,n:954 | 42.09 |
| 43 | Hcri_10_Ane111_ | *Heteractis crispa* | *Amphiprioan clarkii* | *Amphiprion perderaion* | Ourabay, Okinawa | SRR23908449 | 66.4 | 10029 | 173548 | 1917 | 43276 | C:92.2%[S:52.1%,D:40.1%],F:1.7%,M:6.1%,n:954 | 44.68 |
| 44 | Hcri_5_S16 | *Heteractis crispa* | *Amphiprioan clarkii* | | Ankiba, Kagoshima | SRR23908447 | 53.2 | 8026 | 146372 | 1955 | 37800 | C:85.5%[S:62.6%,D:22.9%],F:3.6%,M:10.9%,n:954 | 47.45 |
| 45 | Hcri_6_S17 | *Heteractis crispa* | *Amphiprioan clarkii* | *Amphiprion perderaion* | Oshima District, Kagoshima | SRR23908446 | 45.5 | 6871 | 143255 | 1822 | 33060 | C:89.8%[S:56.8%,D:33.0%],F:2.4%,M:7.8%,n:954 | 43.93 |
| 46 | Hcri_7_S18 | *Heteractis crispa* | *Amphiprioan clarkii* | | Ishigaki, Okinawa | SRR23908445 | 48.3 | 7295 | 145116 | 1968 | 26687 | C:87.5%[S:61.8%,D:25.7%],F:2.9%,M:9.6%,n:954 | 46.92 |
| 47 | Hcri_9_Ane108_ | *Heteractis crispa* | *Amphiprioan clarkii* | *Amphiprion perderaion* | Henza, Okinawa | SRR23908443 | 51.6 | 7797 | 163541 | 1967 | 35302 | C:80.0%[S:59.2%,D:20.8%],F:8.4%,M:11.6%,n:954 | 45.37 |
| 48 | Hcri_8_S19 | *Heteractis crispa* | *Amphiprion clarkii* | | Onnna, Okinawa | SRR23908444 | 39.7 | 5991 | 126,553 | 1,594 | 29259 | C:87.3%[S:67.6%,D:19.7%],F:5.3%,M:7.4%,n:954 | 45.2 |
| 49 | Hcri_2 | *Heteractis crispa* | *Amphiprion clarkii* | | Yomitan, Okinawa | SRR14307522 | 138.7 | 20,949 | 252,997 | 1,986 | 32,026 | C:92.0%[S:53.7%,D:38.3%],F:2.5%,M:5.5%,n:954 | 45.6 |
| 50 | Hcri_1 | *Heteractis crispa* | *A. clarkii, Aperideraion_Dtrimaculatus_gobbi* | | Ishigaki, Okinawa | SRR14307532 | 161.4 | 24,377 | 243,055 | 1,944 | 32,920 | C:80.0%[S:59.2%,D:20.8%],F:8.4%,M:11.6%,n:954 | 45.57 |
| 51 | Haur_3_S20 | *Heteractis aurora* | *Amphiprion clarkii* | | Onna, Okinawa | SRR21497997 | 53.3 | 8,048 | 74,685 | 1,931 | 27,802 | C:88.9%[S:59.0%,D:29.9%],F:4.0%,M:7.1%,n:954 | 40.84 |
| 52 | Haur_4_S1 | *Heteractis aurora* | *Amphiprion clarkii* | | Onna, Okinawa | SRR21497996 | 47.4 | 7,151 | 75,810 | 2,043 | 30554 | C:89.9%[S:60.9%,D:29.0%],F:2.9%,M:7.2%,n:954 | 40.73 |
| 53 | Haur_5_S3 | *Heteractis aurora* | *Amphiprion clarkii* | | Onna, Okinawa | SRR21497995 | 49.4 | 7,455 | 83,496 | 2,016 | 33058 | C:90.7%[S:58.1%,D:32.6%],F:2.6%,M:6.7%,n:954 | 40.67 |
| 54 | Haur_1 | *Heteractis aurora* | *Amphiprion clarkii* | | Yomitan, Okinawa | SRR14307531 | 161.4 | 24,377 | 243,055 | 1,944 | 32,920 | C:92.2%[S:57.4%,D:34.8%],F:1.7%,M:6.1%,n:954 | 45 |
| 55 | Haur_2 | *Heteractis aurora* | *Amphiprion clarkii* | | Yomitan, Okinawa | SRR14307521 | 41.1 | 5220 | 120,341 | 1,721 | 31875 | C:87.6%[S:54.0%,D:33.6%],F:5.0%,M:7.4%,n:954 | 44.1 |
| 56 | N/A | *Cryptodendrum adhaesivum* | N/A | | Australia | SRR14115233 | | | | | | | |
| 57 | N/A | *Macrodactyla doreensis* | N/A | | Australia | SRR14115222 | | | | | | | |
| 58 | N/A | *Macrodactyla doreensis* | N/A | | Australia | SRR14115224 | | | | | | | |
| 59 | N/A | *Exaiptasia* | N/A | | | http://aiptasia.reefgenomics.org/download/aiptasia_genome.proteins.fa.gz | | | | | | | |
